# Coral reef ecosystem functions in a human-dominated world

**DOI:** 10.64898/2026.04.13.712063

**Authors:** Valeriano Parravicini, Michael McWilliam, Nina MD Schiettekatte, Jérémy Carlot, Renato A Morais, Diego R Barneche, Rucha Karkarey, Mehdi Adjeroud, Deron E Burkepile, Jordan M Casey, Maria Dornelas, Graham J Edgar, Dan A Exton, Nicholas AJ Graham, Sally A Keith, Joshua S Madin, Eva Maire, David Mouillot, Nicolas Mouquet, Rick D Stuart-Smith, Giovanni Strona, Sébastien Villéger, Shaun K Wilson, Simon J Brandl

## Abstract

The metabolic processes sustaining coral reefs, from carbonate and primary production to secondary production, remain poorly integrated and rarely quantified simultaneously at global scales. This hampers our ability to predict global responses to accelerating human pressures and manage coral reef functioning. Using metabolic scaling and bioenergetic models applied to surveys from 1,100 reefs worldwide, we provide a global, standardized quantification of 14 ecosystem functions spanning benthic (corals and algae) and fish communities. Our analysis reveals a continuous functional spectrum of global coral reefs organized along four dominant axes: 1) primary production, 2) calcification and habitat structure, 3) secondary biomass production and consumption, and 4) biomass turnover. Functions mediated by fish and benthic communities show weak associations at the global scale rather than tight coupling. Climate stressors reduced calcification and local human impacts lowered secondary production. Yet these directional effects unfolded against a backdrop of substantial natural variability in reef functional configurations, such that heavily and minimally impacted reefs overlap substantially in the global functional space. Temporal analyses across three representative reef systems further revealed that functional trajectories following disturbance are context-dependent, with no universal pattern of recovery across locations. This continuous and context-dependent functional spectrum challenges the notion of universal functional benchmarks and supports locally tailored conservation strategies.

## Introduction

The rates at which ecosystems assimilate, store, cycle, and transfer energy determine their capacity to sustain biodiversity, support human societies, and buffer the consequences of global change^1–3^. For this reason, identifying the major axes of ecosystem functioning and their determinants has become a central goal of macroecology. The diversity of functions in an ecosystem is not random but constrained by climate, community structure, and evolutionary history^4–6^. Analyses of plant functional traits and energy fluxes across terrestrial biomes, for example, have revealed coherent spectra of ecosystem function, providing a foundation for predicting how these systems respond to global change^4^. Yet these frameworks have been developed primarily for terrestrial ecosystems, where vascular plants dominate biomass and simultaneously drive habitat structure, primary production, and carbon storage, leading to strong covariance among functions. In contrast, many marine ecosystems are structured by the joint but potentially independent contributions of phylogenetically and functionally distinct communities. For example, benthic sessile organisms are responsible for the bulk of carbonate production, whereas mobile vertebrate consumers exhibit greater contribution to biomass storage. Whether coherent functional spectra emerge from such systems, and whether the coupling among these assemblages is detectable when observed at different temporal and spatial scales, remains an open question with direct implications for conservation and management.

Coral reefs offer an exceptional opportunity to explore global variation in ecosystem functions. These ecosystems support diverse assemblages whose contributions to reef functioning differ widely: habitat-forming benthic organisms drive calcification, primary production, and three-dimensional habitat structure, while mobile consumers mediate secondary production, nutrient cycling, and trophic transfer^7^. Coral reefs also span all tropical ocean basins and encompass steep gradients of exposure to thermal stress and human impact^8–10^, providing the environmental variation needed to test how functional configurations respond to anthropogenic drivers. Yet their complexity has historically been reduced to analyses of a few broad functional groups, implicitly treating all species within a group as functionally equivalent and generating powerful but potentially oversimplified paradigms^11–13^. While trait-based approaches have begun to decompose this complexity into tractable components^14^, a simultaneous quantification of benthic and fish functions at global scale has not been attempted, partly because the data required have only recently become available. Whether these two community types covary or operate independently therefore remains unknown, limiting our ability to identify the major axes of reef ecosystem functioning and predict how reefs respond to global change.

Here, we use metabolic scaling and bioenergetic models grounded in empirically measured physiological processes to quantify 14 ecosystem functions spanning benthic and fish assemblages across 1,100 coral reefs worldwide. The selected functions capture major metabolic processes and structural features operating in reef ecosystems: for benthic communities, these include gross primary production, organic growth, inorganic growth, calcification, and gross carbon storage as rates of energy and material flux derived from metabolic scaling, alongside rugosity and branch spacing as indices of the three-dimensional habitat structure that emerges from calcification processes and benthic community composition. For fish communities, the functions considered include herbivory, planktivory, piscivory, biomass production, biomass turnover, nitrogen excretion, and phosphorus excretion, all estimated through bioenergetic modelling. Together, this suite of functions spans the principal pathways of energy and material transfer in reef ecosystems, while also capturing the structural dimension of reef functioning that metabolic rates alone cannot reflect, across systems experiencing some of the most rapid environmental change on Earth^7,15^.

## Results

In our global analyses, we first examine correlations between different functions, charting the global spectrum of reef functioning and examining correlations among functions to identify the major axes of reef functional variation. Second, we measure how the functional configuration of reefs changes across space and stress gradients, including thermal anomalies and human impact metrics. Third, we analyze 38 time-series from three Indo-Pacific reef systems to explore the temporal variation in coral reef functions across known cycles of disturbance and recovery.

### The functional spectrum of coral reefs worldwide

Similar to terrestrial ecosystems^4^, we observed high variability in coral reef functions at a global scale, most of which span three to five orders of magnitude across study sites (Fig. 1). For example, reef-scale calcification rates ranged from 0 g CaCO3 m^-2^ hr^-1^ (coral deprived reefs in Costa Rica and Australia) to 12 g CaCO3 m^-2^ hr^-1^ (Diamond Island in the Central Coral Sea, Australia), while secondary consumption rates ranged from 0.05 g C transect^-2^ day^-1^ (O’ahu, Hawaii) to 584 g C transect^-2^ day^-1^ (Malpelo, Colombia). The exceptions were coral reef rugosity and fish biomass turnover, which varied five-fold and one order of magnitude, respectively. Our analysis encompasses calcifying reefs with high to low coral cover, as well as non-calcifying reef systems that are often presented as alternate stable states (i.e., coral-algae phase shifts)^10,16–18^. However, the unimodal distributions across all functions and ocean basins show a continuity of values with high frequencies at intermediate levels (Fig. 1). Indeed, although the presence of zero values in some functions (e.g., calcification) may indicate potentially irreversible shifts, most reefs exist along a continuum, rather than a clustered distribution. This result is further supported by the multidimensional decomposition of functions through a Principal Component Analysis (PCA, see Methods, Fig. 2), which does not yield distinct clusters of reefs, either globally or across ocean realms (Fig. S1). Our results do not refute the existence of alternative stable states but emphasize the vast array of possible functional configurations that cannot be defined by a clear dichotomy between coral and algae-dominated states.

**Figure 1.**
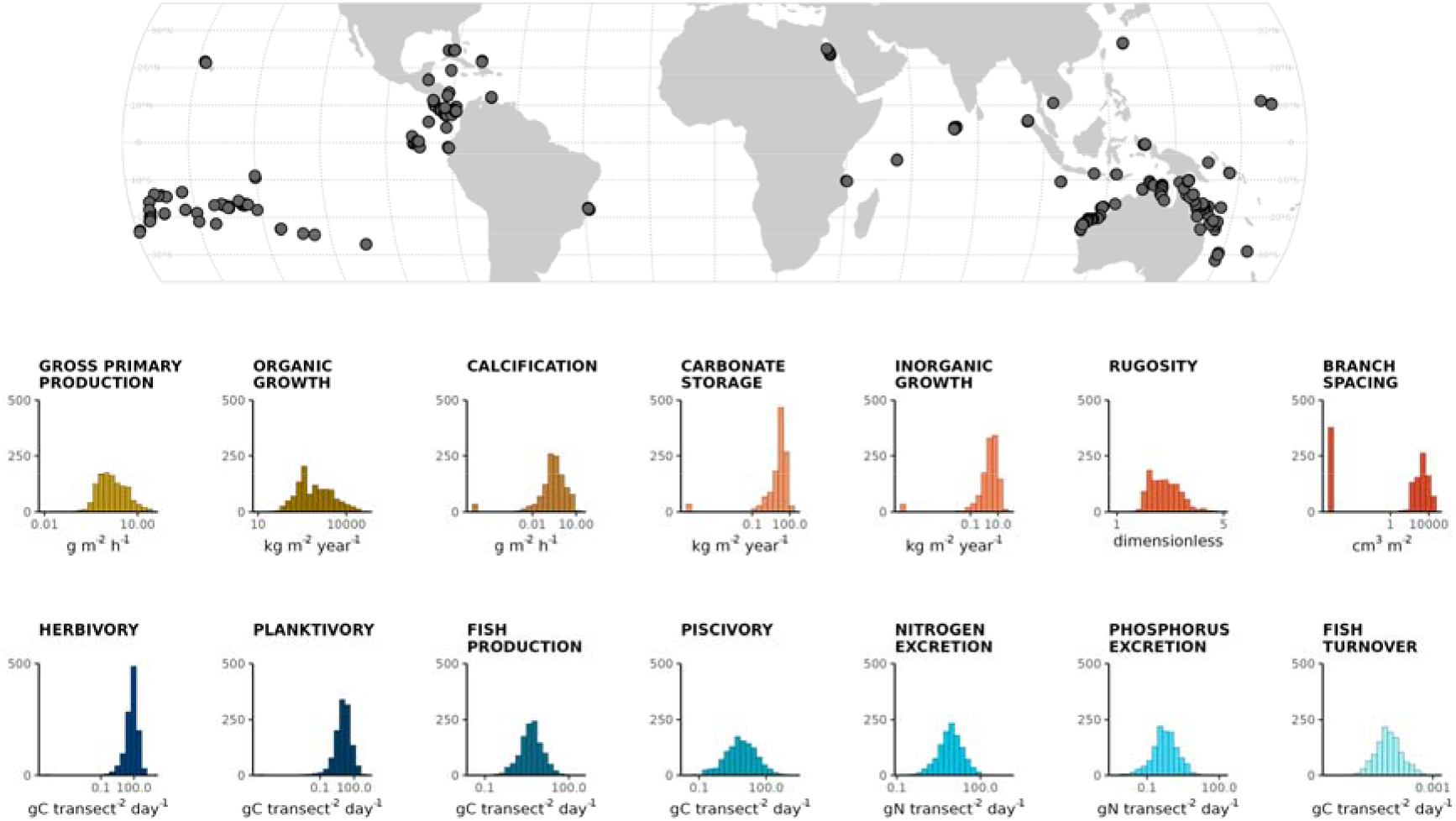
Variability in 14 ecosystem functions across 1,100 circumtropical reefs. Histograms show the frequency distribution of each function across surveyed reefs. Brown histograms represent seven benthic functions: gross primary production (GPP), organic biomass growth, calcification, carbonate storage, inorganic carbonate growth, habitat rugosity, and branch spacing. Blue histograms represent seven fish-mediated functions: herbivory, planktivory, biomass production, piscivory, nitrogen excretion, phosphorus excretion, and biomass turnover. Fish functions were estimated per transect (250 m^2^).

**Figure 2.**
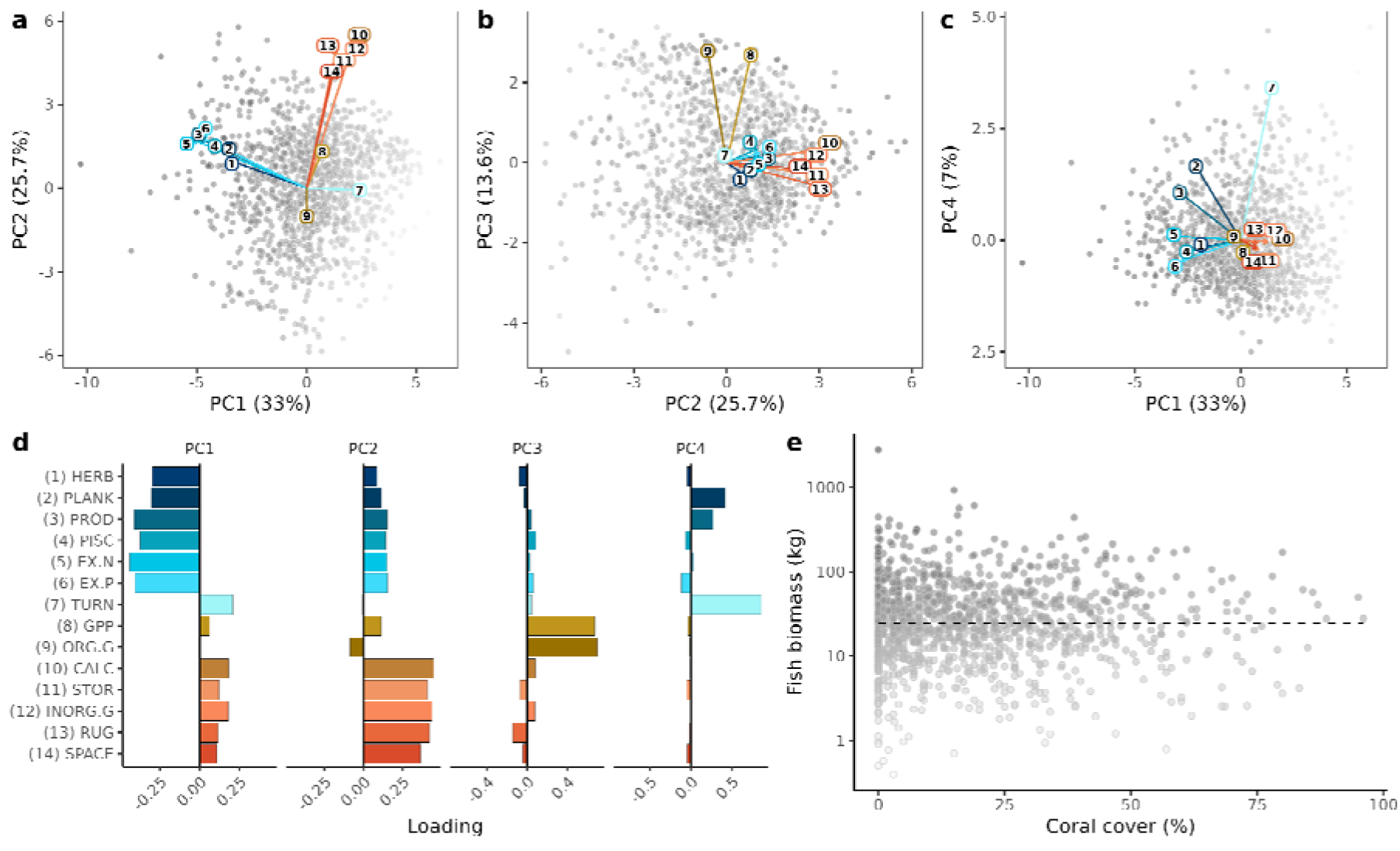
Core functional axes of reefs captured by a principal component analysis (PCA) of 14 benthic and fish functions across 1100 reefs. Panels (a-c) highlight relationships between PCA axes in multivariate space, while panel (d) shows the loadings of the 14 functions onto each axis. We highlight relationships between fish (blue arrows 1-6) and calcifier (red arrows 10-14) functions in PC1 and PC2 (a), between calcifier (red arrows 10-14) and producer brown arrows (8-9) functions in PC2 and PC3 (b), and between fish biomass production (blue arrow 3) and turnover (blue arrow 7) in PC1 and PC4 (c). The relationship between fish biomass and coral cover (e) highlights two principal drivers of ecosystem functions.

To examine relationships between functions across reef ecosystems, we quantified correlations between the primary PCA axes and their loadings (i.e. the contribution of each variable to the principal axes). We found that the multidimensional functional space for coral reefs can be collapsed into four main axes that describe 79.3% of the global variation (Fig. 2 & Fig. S2). The first axis was related to all rates of secondary consumption performed by reef fishes, but not their biomass turnover (PC1, 33% of variation). The second, orthogonal axis was associated with benthic calcification, inorganic growth, and three-dimensional habitat provision (PC2, 25.7% of variation), highlighting the role of calcifying organisms in creating long-lasting, architecturally complex frameworks. The third axis (13.6% of variation) describes functions performed by primary producers (specifically gross primary productivity and organic growth), while the fourth axis (7% of variation) is characterized by functions performed primarily by small fish species, specifically biomass turnover and, to a lesser degree, planktivory. These results reveal that reef ecosystems worldwide commonly support multiple functions operating at high rates while showing low rates across other functions, likely due to the overwhelming role of fish biomass and coral or algal cover (Fig. 2d). This corroborates global functional patterns in terrestrial systems, where the primary PCA axis is driven by collinearity among multiple functions (e.g., productivity, respiration, evapotranspiration) related to increasing plant biomass across ecosystems^4^.

Importantly, despite the widely assumed strong relationships between benthic and fish communities, we found largely orthogonal relationships between functions performed by the two communities (Fig. 2a & Fig. S3), which is also highlighted by the lack of correlations between fish biomass and coral cover (Fig. 2e). For example, we found that calcifiers functions (i.e. calcification and carbon storage) and complexity metrics (i.e. rugosity and branch spacing) are largely uncorrelated to fish functions, although planktivory shows a weakly positive relationship with those metrics (Fig. S3). However, while carbonate production is predominantly driven by fast growing corals, which also create structures that support fish biomass and productivity^19^, the structural complexity provided by live coral, may be replaced by coral skeletons and reef geomorphology or other benthic biota, such as soft corals, sponges, or canopy forming macroalgae, enabling the maintenance of high fish productivity^20^. As a consequence, relationships between fish and benthic productivity from different reef types may be obscured when data are viewed globally. Moreover, the links between fish and coral functions may be further blurred as benthic communities change and the composition of fish assemblages becomes more reliant on the benthic structures that replace coral. Anyway, the few links between benthic and fish-mediated functions provide further support to recent evidence of weak correlations between live coral cover and fish species richness or abundance^21–23^. Furthermore, we did not identified strong negative relationships within each component (Fig. S3). For example, while we may have expected a negative relationship between carbonate production and organic production (assuming the first is dominated by corals^24^, the second by algae^25^), the occurrence of such expected trade-offs is likely buffered by numerous reefs with simultaneously low cover of corals and algae. The relationship does become apparent when higher values of coral and algal cover are considered, suggesting these and other relationships between functions may be non-linear and that ecological thresholds may exist (Fig. S7).

### Human impacts on reef functions

The heat-sensitivity of corals makes coral reefs one of Earth’s most vulnerable ecosystems. Therefore, we sought to quantify the impact of persistent heat stress (using Degree Heating Weeks the year before the survey, DHW) and local anthropogenic pressures (using human gravity^26^ and protection status) on the global variation in reef ecosystem functions (see Methods). Our analysis revealed broadly expected responses of the functional spectrum of coral reefs exposed to these anthropogenic impacts (Fig. 3). Specifically, DHW was positively correlated with functions related to primary producers (e.g., gross primary production and organic growth) that typically increase in abundance following coral bleaching and mortality. Human Gravity was negatively correlated with all fish functions except biomass turnover, primarily leading to decreases in consumption rates (piscivory and herbivory) and nitrogen excretion (Fig. 3). A weak negative association emerged between human gravity and functional axes associated with calcifier functions, which is also evident when individual functions are analyzed separately (Fig. S4). No-take marine protected areas were associated with a general increase in fish functions (mostly piscivory and planktivory), but did not positively influence benthic functions (Fig. 3 & Fig. S4). These patterns should be interpreted in light of the inherent limitations of global-scale analyses: aggregate relationships across thousands of reefs may obscure locally consistent (and relevant) but regionally heterogeneous effects, an ecological form of Simpson’s paradox in which a trend observed at the global scale reverses or disappears when examined within specific regions or reef systems^21,22^. Indeed, while the effects of anthropogenic drivers on coral cover are detectable at the global scale, their consequences for benthic community composition, a key driver of functions, are considerably weaker at this scale^27^, suggesting that local and regional analyses remain essential complements to the global perspective offered here.

**Figure 3.**
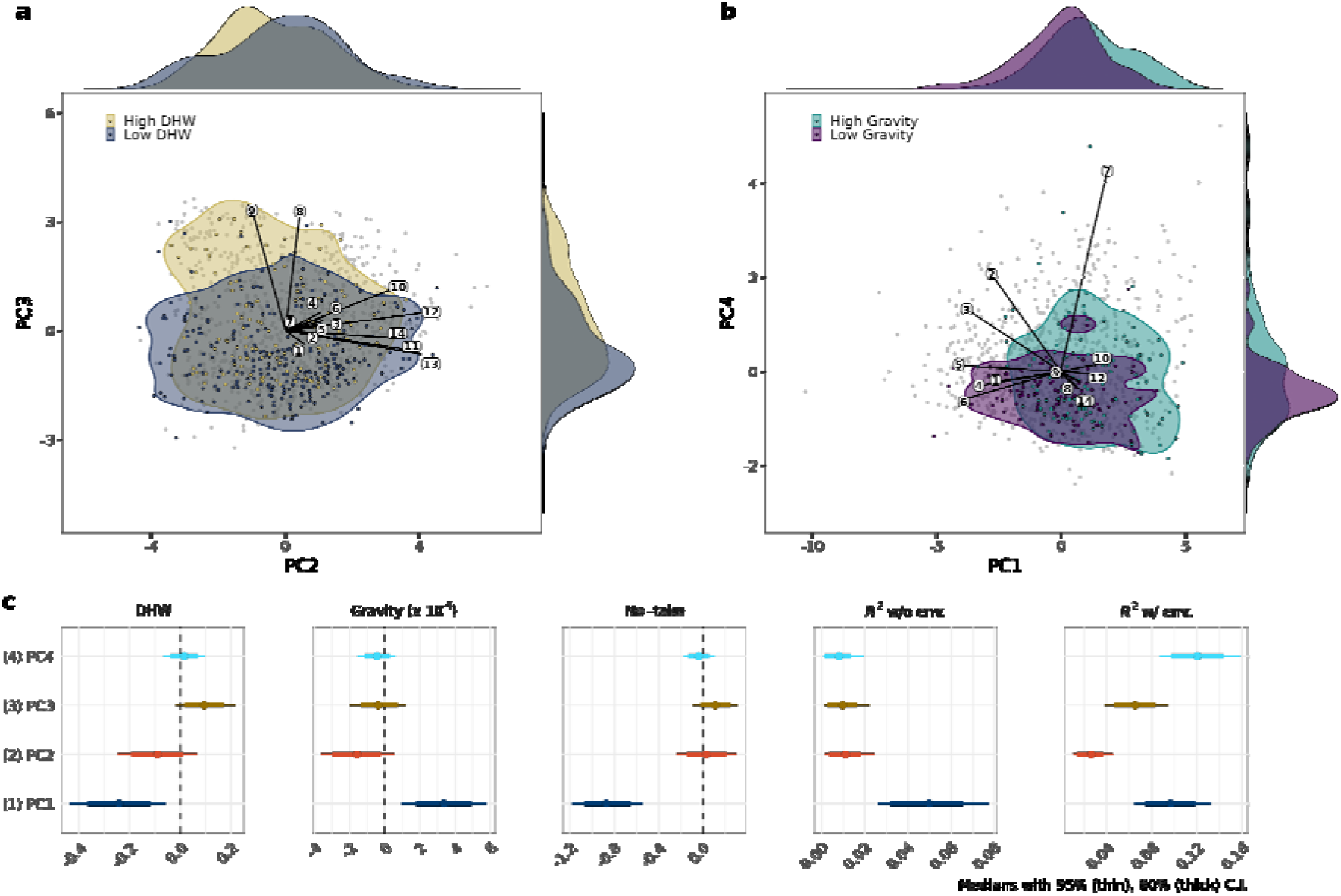
Human impacts on reef functions in multidimensional space. (a) Shifts along the functional axes across reefs exposed to the upper and lower 5% of thermal exposures (quantified by degree heating weeks, DHW). Only PC2 and PC3 are presented due to their role in calcifier and producer functions, respectively. (b) Shifts along the functional axes across reefs exposed to the upper and lower 5% of human gravity estimates. Only PC1 and PC4 are presented due to their role in fish biomass production and turnover, respectively. (c) Summary of models linking human stressors (DHW, human gravity and no-take status) to the four functional dimensions. The x-axis indicates the effect size derived from a Bayesian hierarchical model. PC1 is related to consumption and secondary production, PC2 to calcification and habitat structure, PC3 to primary production and PC4 to secondary biomass turnover.

Nevertheless, our models highlight the independent nature of the footprint of climate change and human proximity on coral reef functioning^28^. Specifically, human gravity and DHW related to the spectrum of reef functions following two orthogonal axes. In particular, DHW and local stressors move the functional spectrum toward functions dominated by high carbon turnover, limited carbonate storage, and decreased habitat provision, a pattern that has likewise been observed in the terrestrial realm where large trees that provide structure and carbon storage are replaced by smaller and more ephemeral species^29^. However, importantly, models including these human footprint variables explained less than 15% of the global variation in reef functions when additional environmental variables were included, and even less on their own (Fig. 3). In the PCA space of reef functions, the least impacted reefs (comprising the 5% of reefs with the lowest DHW or human gravity) delineate a polygon that broadly overlaps with the most heavily impacted reefs (comprising the 5% of reefs with the highest DHW or human gravity, Fig 3a,b), raising questions over the predictive power of these anthropogenic variables for reef functional configurations. While there are negative effects of warming and local human pressure on coral reefs, reefs occupy a vast spectrum of functional configurations even under considerable anthropogenic stress. This finding suggests that functionally impacted reefs retain functional capacity and associated ecosystem services, although the species that provide these functions and services may change^30^.

### Reef functions following disturbance and recovery

To deepen our analyses of reef functional configurations in the face of disturbance, we investigated temporal shifts in fish and benthic functions in time series from 38 sites across three representative Indo-Pacific reef systems (Seychelles, Indonesia and French Polynesia), each with known histories of disturbance-recovery cycles. Our results show broadly diverging shifts in coral reef functions across the three case studies, even across sites that are separated by only a few kilometers (Fig. 4b). The Seychelles experienced severe impacts from global coral bleaching in 1998^31^, leading to consistent decreases in fish turnover and calcifier functions (i.e. carbon storage, calcification and rugosity)^32^ but starkly divergent trajectories in organic production (Fig. 4b). Sites in French Polynesia were subjected to substantial reductions in coral cover due to an outbreak of crown-of-thorns starfish (*Acanthaster planci*) followed by a severe tropical cyclone^33–35^. This disturbance was associated with a decline and recovery of calcifier functions, but highly variable trends in primary producer functions, perhaps due to differences in hydrodynamics across sites^36,37^. At the same time, consumption and secondary biomass production tended to decrease. In Indonesia, reefs were impacted by bleaching in 2010 and 2016^38^, yet they displayed high stability in benthic functions, possibly because of the relatively small change in coral cover from 30% to 20% (Fig. S5). At this location, there was an overall decrease in secondary biomass production (Fig. 4). If the expected negative relationships between disturbance and coral reef functions were consistent, predictable or inevitable, we would expect them to emerge every single time. The fact that we found as many instances of negative, positive or negligible relationships in just 38 sites across 3 regions illustrates the high context-dependency and regional variability in ecosystem function trajectories following acute disturbance. Such context-dependency may reflect the influence of legacy effects, including the duration and intensity of prior disturbance and the historical composition of local communities, which modulate how reefs respond functionally to acute events^39^.

**Figure 4.**
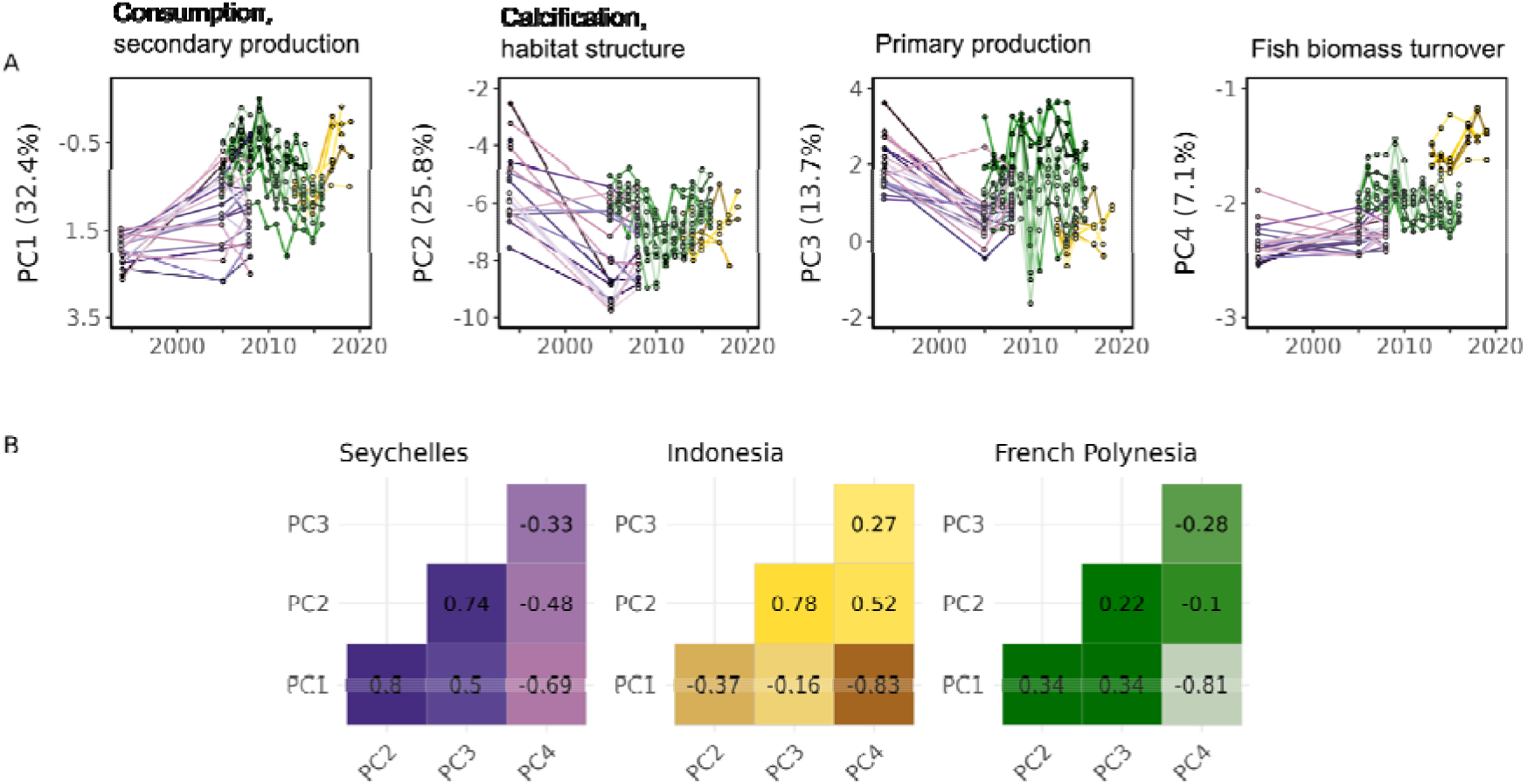
Changes in reef functions over time across three Indo-Pacific reef systems. (Seychelles in purple, Indonesia in yellow and French Polynesia in green). Upper panels display the transitions through functional space for reefs in each location through time. Lower panels show correlation coefficients for the functional axes over time for all three Indo-Pacific reef systems.

Our temporal analysis also revealed the importance of spatial heterogeneity for the interdependence between the functional axes over time. Specifically, while there were some strong correlations between different functions (including benthic and fish-mediated functions) at the local scale (i.e. within a location), there was no overarching pattern across the Indo-Pacific in the co-variation of different functional axes. For example, all fish functions, except fish Biomass Turnover, were strongly positively correlated with calcifier functions in the Seychelles. Yet, this was not observed in French Polynesia or Indonesia, where negligible or negative relationships between PC1 and PC2 were observed, respectively. Therefore, while our temporal analysis uncovered limited associations between benthic and fish functions, such connections were clearly dependent on the nature of the disturbance and regional or local conditions. These divergent trajectories further confirm that functional coupling between benthic and fish communities is highly context-dependent.

## Discussion

By projecting organism metabolism onto community-level data, we reveal the major axes of coral reef ecosystem functioning at a global scale, showing substantial variation across space, time, and perturbation gradients. Climate stressors and local human impacts drive directional shifts in reef functional configurations, reducing calcification and secondary production in ways consistent with documented trends in reef degradation. Yet these effects unfold against a backdrop of enormous natural variability, such that anthropogenic drivers alone explain a modest fraction of the global variation in reef functions. This does not diminish the conservation urgency of addressing climate change and overfishing, the effects of which are clear and directional and will continue to challenge reef conservation and management^40^. However, it does suggest that reefs may retain a broader functional repertoire under stress than is often assumed, with implications for how we define management targets. Indeed, our results provided no evidence for a universal ‘pristine’ functional state that could serve as a global benchmark. Instead, our analyses revealed a large and continuous array of functional configurations across reefs exposed to a wide range of thermal and fishing stresses (Fig. 3). Locally tailored conservation strategies are needed to account for the functional context of individual reefs, rather than the pursuit of a single idealized state.

The functional independence of benthic and fish communities likely reflects their contrasting responses to drivers operating at different temporal and spatial scales. Benthic communities are primarily shaped by thermal disturbance and physical factors driving slow dynamics of coral growth and calcification, while fish assemblages respond more rapidly to fishing pressure and habitat availability. At the global scale, where these drivers vary largely independently, functions performed by the two communities are decoupled, consistent with recent evidence of weak correlations between coral cover and fish abundance across broad spatial^41^ and temporal scales^42^. The decoupling of fish and benthic functions apparent in global analyses may be context dependent and an emergent outcome of multiple stressors or processes acting on communities with fundamentally different response timescales.

Recognizing both the directional effects of anthropogenic stressors and the context-dependency of benthic-fish interactions, our results support the need for local and national efforts for the conservation and management of coral reefs^43^, acknowledging the likelihood of ongoing changes and potentially impossible task of reverting to historical conditions^44,45^. For instance, our temporal analyses support a multitude of local studies showing that increased macroalgal production at the expense of scleractinian corals^18,30,46^, a generally undesired benthic transition, may be temporally associated with increases in several fish functions (such as biomass production by herbivorous fishes) in certain locations (Fig. 4). In fact, despite the general lack of correlations among fish and benthic functions at the global scale, our results do not rule out the emergence of such correlations at local, short-term scales. Rather, our results show that high secondary production, for example, may be found on reefs characterized by both high coral and algal cover, which may offer pathways for sustaining important contributions to people in the face of heat stress and benthic degradation. This pattern is consistent with top-down and bottom-up regulation mechanisms, whereby high fish biomass can be sustained by both coral-dominated and algae-dominated benthic states depending on the availability of prey and habitat structure^17,18^. Similarly, elevated herbivory rates may be present on reefs dominated by macroalgae, as well as on reefs with high coral and algal turf cover^47^. The large variability of coral reef functions calls for conservation strategies that maximize ecological integrity and human wellbeing, rather than a pursuit of a narrow range of idealized states that do not exist or cannot be defined through a single functional configuration.

Finally, our approach enables a better understanding of coral reef functioning across spatial and temporal gradients, and facilitates functional comparisons of ecosystems that would not be possible through compositional analysis alone^11^. Our methodology builds upon extensive literature refining approaches to scale individual traits to estimate community processes^34,48–52^ and has been validated with *in situ* estimation of fish excretion^50^ and gross primary production (Fig. S12). While our scaling approach cannot (and should not) replace direct measurements of biogeochemical fluxes^53^, it provides a foundation to estimate variability in the transfer of energy and nutrients across reefs worldwide and its underlying drivers. For example, our analysis uncovers the tendency of human-impacted reefs to express higher rates of functions associated with high carbon turnover and limited storage, which corroborates concerning trends in other ecosystems^29^. Beyond coral reefs, this has implications for large scale nutrient fluxes (especially for carbonates, for which reefs play a globally dominant role) and directly addresses processes of inherent human interest, such as fisheries productivity^54^. The enhanced capacity to gauge changes in ecosystem dynamics across the revealed functional spectrum is essential to understand local reef context and implement effective measures to protect our planet’s ecosystems in response to rapidly unfolding global changes.

## Materials and Methods

### Survey Data

To determine spatial and temporal trends in coral reefs, we used the Reef Life Survey dataset (RLS)^55^ and a compilation of 38 time series datasets across three locations: Moorea (French Polynesia), Hoga (Indonesia) and the inner Seychelles (Fig. 1). For the spatial analysis we used RLS underwater visual surveys conducted between September 2006 and May 2019, which include both fish and benthos assessments at the same sites. In the present work, following Parravicini et al. (2013)^56^, we employed a broad definition of tropical coral reefs, focusing on locations with annual average Sea Surface Temperature (SST) higher than 17°C. Fish surveys are conducted along 50m-long and 5m-wide belt transects. From the RLS data, we excluded cryptobenthic fishes^57^ and elasmobranchs because their biomass is not appropriately assessed through the standard visual surveys and small transects, respectively. Mobile invertebrates were similarly excluded as their abundance and biomass cannot be reliably quantified through the visual census methodology. While these groups contribute to reef functioning, their exclusion is consistent with standard practice in large-scale reef monitoring and does not affect our estimates of fish and benthic community functions.

Fish abundance and size was then converted to biomass using length-weight conversion coefficients available from Fishbase^58^. Information on the percentage cover of sessile animals and algae is collected along the transect lines set for fish counts using photo-quadrats taken sequentially every 2.5 m along the 50 m transect. Digital photo-quadrats are taken vertically downward to encompass an area of 0.3 m × 0.3 m. The percentage cover of different macroalgal, coral, sponge and other attached invertebrate species in photo-quadrats is digitally quantified on the computer using a grid of 5 points overlaid on each image, with the taxon lying directly below each point being recorded. Each transect, thus, comprises 20 photos, from where benthic cover composition is determine from 100 points per transect. All hard (e.g., scleractinian) and soft (e.g., alcyonarian) corals are identified to the genus level. For survey points that do not intersect live coral, the underlying habitat is categorised as macroalgae (> 5 mm in height; identified to genera), turf algae (< 5 mm in height), crustose coralline algae (CCA), rubble or sand.

For the temporal analysis, we collated a dataset of 38 time series from three locations (Seychelles, Indonesia and French Polynesia, Table S1). These time series met several criteria that used for their selection: data were available and known to the authors, with surveys including both fish and benthic surveys, their taxonomic resolution was comparable to that of the RLS dataset. Time series were intentionally selected to encompass one complete disturbance-recovery cycle rather than to maximize temporal coverage, as our aim was to characterize functional trajectories following acute disturbance events rather than long-term trends.

From French Polynesia, we used data from 11 sites around Mo’orea obtained each year from 2005 to 2015. In 2005 the reef was relatively undisturbed and coral cover was at the historical maximum (∼40-50%) Then, the reef was severely impacted by a 2-year outbreak of the Crown-of-Thorns Starfish that started in 2007, and subsequently from severe Cyclone Oli in 2010. Together, these disturbances reduced coral cover down to 2%. Subsequently, coral cover recovered steadily, reaching pre-disturbance values in 2015^59,60^. In this location, fish counts were obtained from 250m^2^ transects, while benthic communities were assessed from 20 1m^2^ photographs. For the Seychelles, we used data previously described by Graham et al.^31^. Briefly, data refer to 21 sites spread across the inner Seychelles, where benthic and fish communities were surveyed in 1994, before the mass coral bleaching of 1998, and then in 2005 and 2008. Twelve sites recovered to pre-disturbance conditions while the others turned into algae-dominated reefs. Fish counts consisted of fixed counts over an area of ∼150m^2^ and the cover of benthic categories was visually estimated from the same area. For Indonesia, we obtained time series at 6 sites around Hoga Island in the Wakatobi National Park, Sulawesi. These sites were surveyed annually between 2013-2015 and again between 2017-2019 before, during and after the global bleaching of 2016 and 2017. Fish counts were obtained from 250m2 transects using diver-operated stereo-video, and benthic communities surveyed along 50m point intercept transects with data points every 0.5m. The reefs around Hoga were largely spared the sudden onset of mass bleaching seen elsewhere during global events, but have suffered heavily from the impacts of intensive artisanal fishing pressure to both the fish and benthic communities^61^.

### Fish Functions

We quantified seven functions performed by fish assemblages: nitrogen excretion rate (gN transect^-2^ day^-1^), phosphorus excretion rate (gP transect^-2^ day^-1^), production of biomass through growth (gC transect^-2^ day^-1^), herbivory (i.e., ingestion rate of macrophytes (gC transect^-2^ day^-1^), piscivory (i.e., ingestion rate of fishes (gC transect^-2^ day^-1^), planktivory (i.e., ingestion rate of plankton (gC transect^-2^ day^-1^) and biomass turnover, defined as the proportion of fish biomass that is expected to be replaced by one year of biomass production (i.e. ratio between biomass production and biomass expressed year^-1^)^18^.

To calculate the seven ecosystem functions carried out by fish communities, we used a bioenergetic modeling approach that predicts rates of physiological processes at the individual-level by taking into account species identity, body size, and temperature^62^. This bioenergetic model is based on metabolic theory and ecological stoichiometry and requires a number of parameters (e.g. growth rate, metabolic rate, diet and body stoichiometry) to estimate elemental fluxes. We used a combination of empirical data and phylogenetic extrapolation to estimate parameters for each species following the methods described in Schiettekatte et al. (2022)^62^. Compared to Schiettekatte et al. (2022), however, we updated previously used constants for absorption efficiency by extrapolating empirical estimate of absorption efficiencies based on species diet and phylogeny (using data from Schiettekatte et al. 2023^63^). We limited our extrapolation only to families for which empirical data were available since we felt confident extrapolating at the genus or species level, but not at family levels. Therefore, we excluded fish families for which we had no parameters, which amounted to <1 % of species across the total dataset. Once individual processes (e.g. growth, consumption, excretion) were accounted for by these predictive models, ecosystem functions were estimated for whole communities by summing the contributions of all individuals. For diet-specific functions (herbivory, piscivory, and planktivory), we summed consumption rates across all individuals classified in each of the associated trophic guilds following Parravicini et al. 2020^64^.

### Benthic functions

We quantified seven benthic functions from monitoring data using scaling relationships to estimate the total production and metabolic activity of each community^65^. First, we used transect-based estimates of percent cover from eleven groups encompassing the major benthic taxa on coral reefs. These included six groups of corals (tabular, branching, corymbose, laminar, hemispherical, and encrusting), four groups of algae (turf algae, fleshy algae, coralline algae, and *Halimeda*), and one substrate category (sand and/or rubble). Normalized estimates (per unit area or volume) of metabolism, stoichiometry, morphology, and growth were estimated from the literature for each group, with coral data derived from the coral trait database^66^ updated with recent literature^67,68^, crustose coralline algae data from Cornwall *et al*.^24^ and data for turf and fleshy algae from an independent literature search (Table S2).

Projections from planar cover to three-dimensional (3D) were used to scale up normalized rate values to the community-level. Given the lack of individual/colony-level data for benthic groups, we estimated the 3D area and volume contributed by each taxonomic group by simulating a size structure for each group while keeping the percent cover values constant. This was followed by projecting the body sizes into 3D ‘morphs’ using data on organism geometry (e.g., height, shape, and branching dimensions, Table S3). Size structure for each community was simulated using two different size distributions: uniform (all the same size, mean diameter equals 10 cm; mostly juveniles), mixed groups (a mix of juveniles and group-averages), and group averages (mean diameter higher than 20 cm; adults). The analysis was performed for each of the three size distributions to test the sensitivity of our results to the simulations (Fig. S5). We conducted a ground-truthing analysis to test whether our estimates of 3D surface area based on organism geometry (Table S3) were similar to the values obtained from 3D laser scans (Fig. S8).

The seven benthic functions were estimated by scaling up metabolic rates by the total area or volume of each taxon (Table S4). Gross photosynthesis and calcification (in g hr^-1^) were measured by projecting metabolic rates per unit area by the total 3D area of each group. Projections of photosynthesis and calcification included an allometric scaling coefficient with 3D area to account for disproportionate changes in body shape with organism size in the main groups^68^. Organic and inorganic production (kg yr^-1^) were estimated as changes in the volume of biomass or carbonate, respectively, following one year of growth in each organism. The standing stock of carbon before modelling growth was used as an estimate of carbonate storage. Organic growth was projected by 3D area for corals and volume for algae. Rugosity was defined using Husband *et al*.^69^, and branch spacing (in cm^3^) was estimated by subtracting organism volume from the 3D convex hull of the organism derived from its geometric attributes (Table S3). We ground truthed our productivity estimates by comparing the distribution and range of productivity values derived from our scaling approach with estimates based on *in situ* metabolic chambers (Figure S11).

Functions were temperature-corrected using the mean (5-year) sea surface temperature at each site. Temperature corrections were conducted separately for corals and algae to account for different thermal tolerances. Corrected coral functions were estimated using the thermal performance curves in Álvarez-Noriega et al.^70^, using the equation:

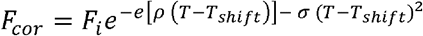

where *F*_*i*_ is the uncorrected function at site *i, T*_*i*_ is the ambient temperature in °C, and to be conservative, *T*_*shift*_, *ρ* and *σ* are constants derived from *A. hyacinthus* in Álvarez-Noriega et al. (30.197, 1.022, and −4.518, respectively). *A. hyacinthus* was selected for this correction because it exhibits the lowest thermal optimum and fastest overall growth among coral species for which empirical thermal performance data are available, providing a conservative estimate of temperature sensitivity across the range of coral communities surveyed. Corrected algal functions were estimated using the thermal performance curves in Terada *et al*. ^71^ using the equation:

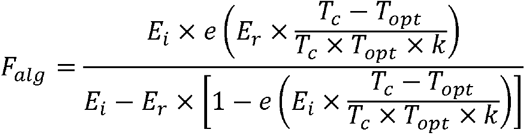

Where *T*_opt_ = 288.15 + plogis(*T*_opt_′) × 30; *E*_r_ = *E*_i_ × plogis(*E*_r_′); and *k* is the Boltzmann constant (8.62 × 10^−5^), *T*_*c*_ is the ambient temperature in Kelvin (K), and T_opt_’, E_r_’ and E_i_ were constants derived from Terada *et al*^71^ (0.21, −1.31, and 3.00, respectively). See Figure S8 for the correction factors.

### Environment-Function Relationships

We used a Bayesian Hierarchical Linear Model to examine the associations between each axis of the functional spectrum and several environmental (Chlorophyll-a, Net Primary Production, Salinity, Sea Surface Temperature, pH) and anthropogenic variables (Human Gravity, Degree Heating Weeks, and the presence of no-take Marine Protected Areas).

Chlorophyll-a (Mg m^-3^) and Salinity (PSU) were obtained from the Copernicus Ocean Data^72,73^. Data were extracted for the periods from 1997 and 2019 and from 1994 to 2017, respectively and annual average data was extracted at the RLS locations from rasters of 0.25° × 0.25° resolution. Sea Surface Temperature (SST, °C) and Degree Heating Weeks (DHW) were extracted by the NOAA CoralWatch v3.1 dataset. Both datasets globally consist of data obtained for the period between 1985 and 2019; we extracted data referring to the period between 1 and 5 years before the exact date of the surveys. In particular, SST was the average of the monthly SST for the 12 or the 56 months before the surveys while DHW was calculated as the accumulation of daily temperature - maximum monthly mean SST from 1985-1993), when this difference exceeded 1°C across windows of a 12 week period (see Liu et al.^74^. Net Primary Production (NPP, mg C m^2^ day^-1^), expressed as the average of monthly data for the period 2003-2018, was extracted from a raster provided by the Oregon State University at a resolution of 0.25°^75^. pH was obtained from a raster dataset provided by the Norwegian Earth System Model forced ocean simulation (NorESM2, https://wiki.met.no/noresm/start). As per SST and DHW, we extracted the mean value for 1-year and 5-years before the date of the surveys. Human gravity represents a composite measure of human influence that combines travel time and human population density, and was obtained from Cinner et al.^76^. Information on the presence of no-take Marine Protected Areas was obtained from the original RLS dataset.

The models were performed in *brms*^77^ at the level of individual transects. The untransformed values of PC axes were used as response variables using a gaussian distribution and weakly informative priors for the slope (a ∼ *normal* (0,1)). The site, as defined in the RLS dataset, was modelled as a random effect for the intercept. Environmental variables (see above), DHW, Human Gravity and the presence/absence of ‘no-take’ MPAs were used as explanatory variables. To address collinearity among environmental variables, we incorporated the first two axes of a separate Principal Component Analysis (PCA) into the model. We ran the models for 5,000 iterations and 2,500 iterations as warm-up. This model structure was run with and without environmental variables in order to compare the explanatory power of climatic and anthropogenic effect vs. local environmental variation. Model convergence was assessed by the visual inspection of traceplots, posterior predictive check plots (Fig. S10 & Fig. S11), and Gelman-Hubin diagnostic values. The same models were also run separately on each of the 14 functions. In this case the response variable was log-transformed and modelled with a gaussian distribution using, as above, weakly informative priors on the slope. Sites were again modelled as a random intercept, and we ran the models for 5,000 iterations and 2,500 iterations as warm-up.

### Temporal analysis

For each site in each location of our temporal dataset (including inner Seychelles, Moorea, French Polynesia and Hoga, Indonesia), we estimated the same seven benthic and fish functions that were also estimated for the spatial dataset, then obtaining their functional space PCA coordinates in by predicting PC scores from the PCA calibrated with the larger RLS dataset. For each location we then explored the relationship between each PC axis using a correlogram.

## Acknowledgments

This research is a product of the “SCORE-Reef” group, supported by the synthesis center CESAB of the French Foundation for Research on Biodiversity (FRB) and the Office Français de la Biodiversité (OFB). We thank Reef Life Survey divers and all data collectors for their contributions to the paper. We thank Peter Esteve for providing valuable support in managing the computational server used to conduct data analyses.

## Notes

### Competing Interest Statement

The authors have declared no competing interest.

