## Supporting information for "Coral reef ecosystem functions in a human-dominated world"

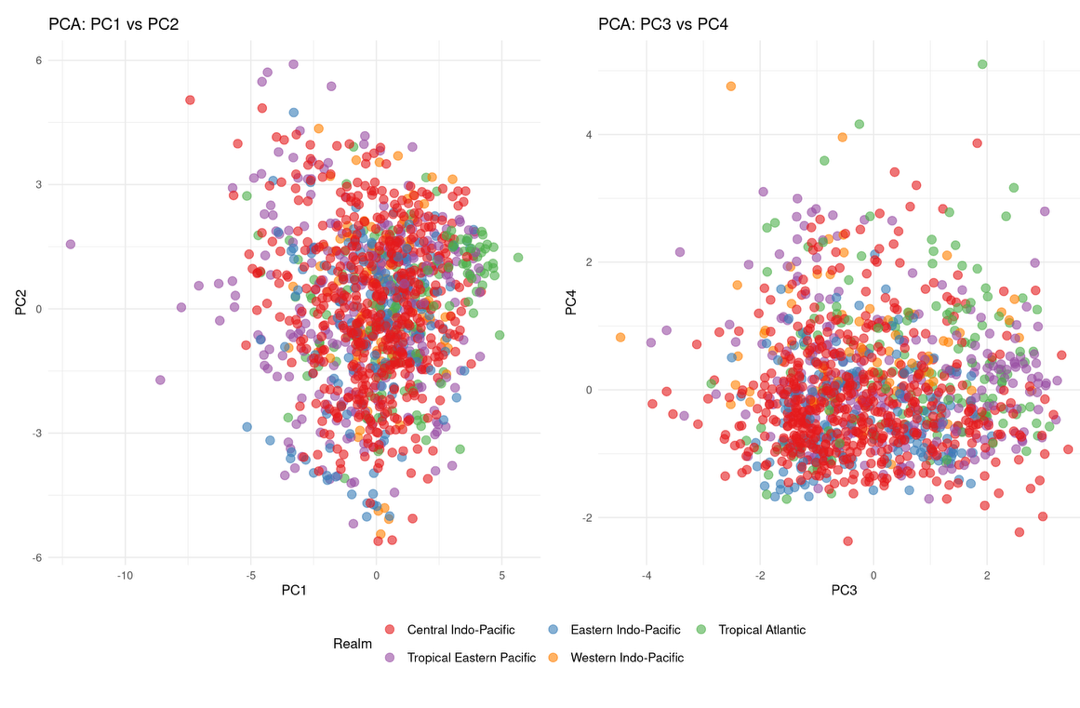

**Figure S1**. Functional space of coral reefs across ocean realms. Principal Component Analysis (PCA) scores for PC1 vs PC2 (left) and PC3 vs PC4 (right) for 1,100 coral reefs worldwide, colored by biogeographic realm. The broad overlap across realms indicates that the continuous functional spectrum documented globally is not an artifact of pooling ecologically distinct regional systems, but reflects genuine variation within and across ocean basins.

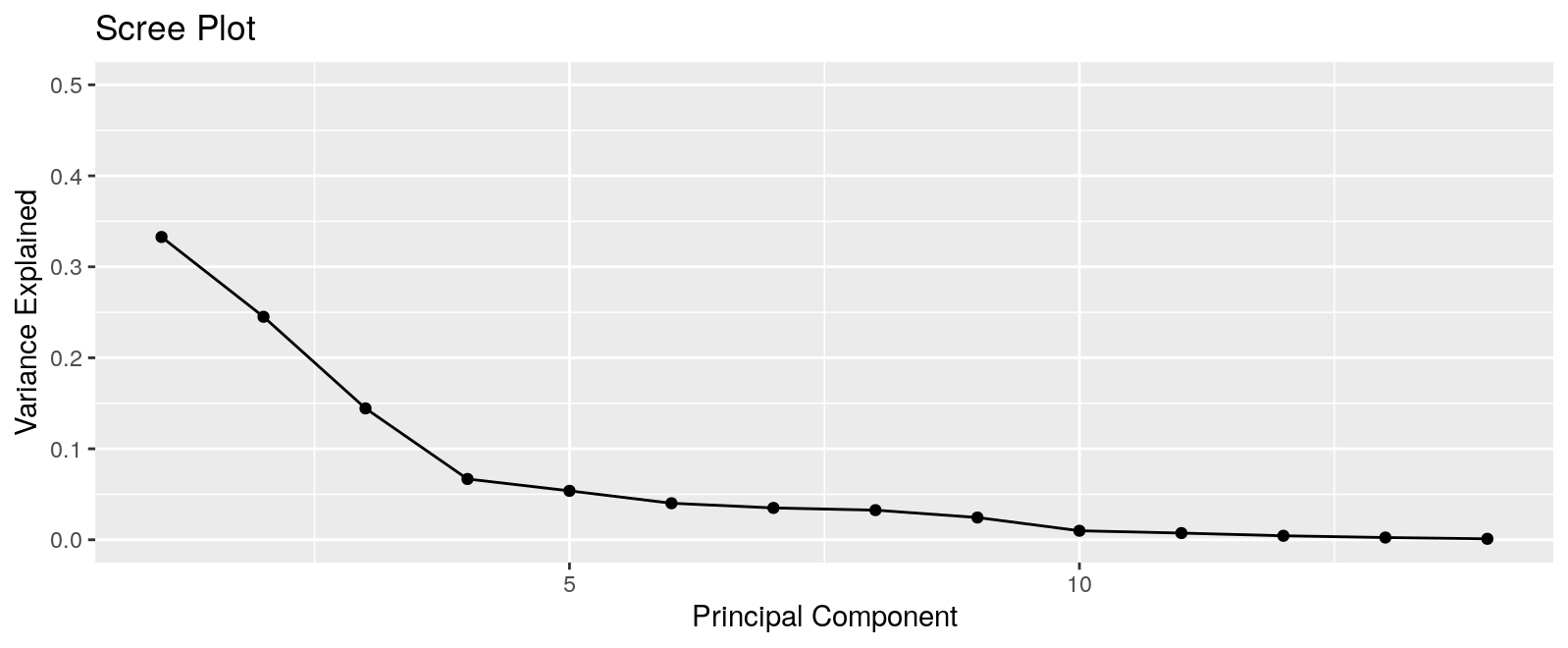

**Figure S2.** Scree plot showing the variance explained by the Principal Components of the PCA on the 14 coral reef functions presented in Figure 2.

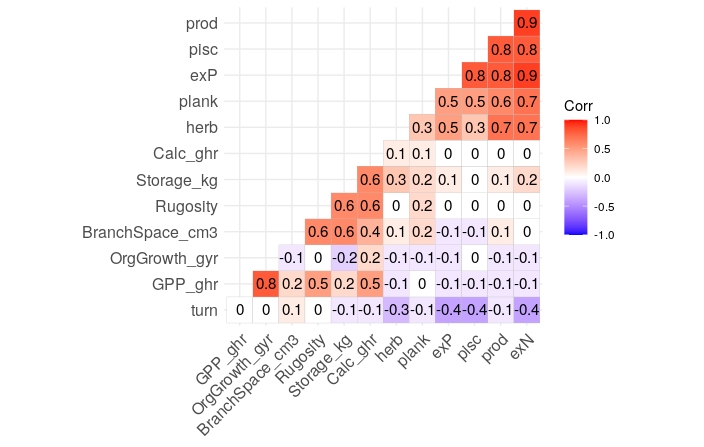

**Figure S3.** Correlogram showing the correlation between the 14 functions estimated for benthic and fish functions including *Herbivory* (herb), *Planktivory* (plank), *Biomass Production* (prod), *Piscivory* (pisc), *Nitrogen Excretion* (exN), *Phosphorus Excretion* (exP), and *Biomass turnover* (turn), Gross *Primary Production* (gpp_ghr), *Organic Biomass Growth* (OrgGrowth_ghr), *Calcification* (Calc_ghr), *Carbonate Storage* (Storage_Kg), *Habitat Rugosity* (Rugosity), and *Habitat Space* (BranchSpace_cm3)

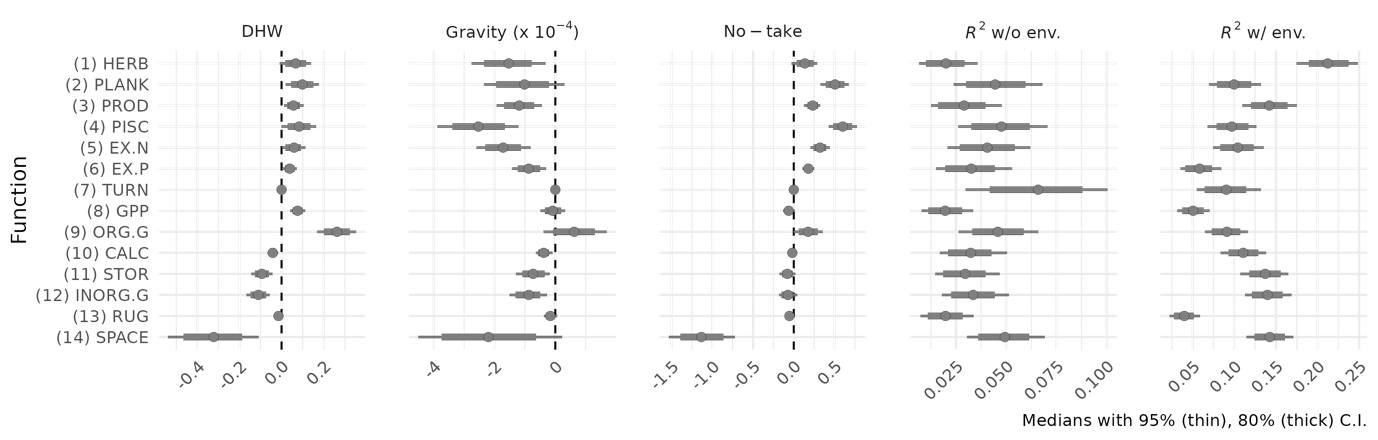

**Figure S4.** Summary of models linking human stressors (DHW, human gravity and no-take status) to the 14 functions. The x-axis indicates the effect size derived from a Bayesian hierarchical model.

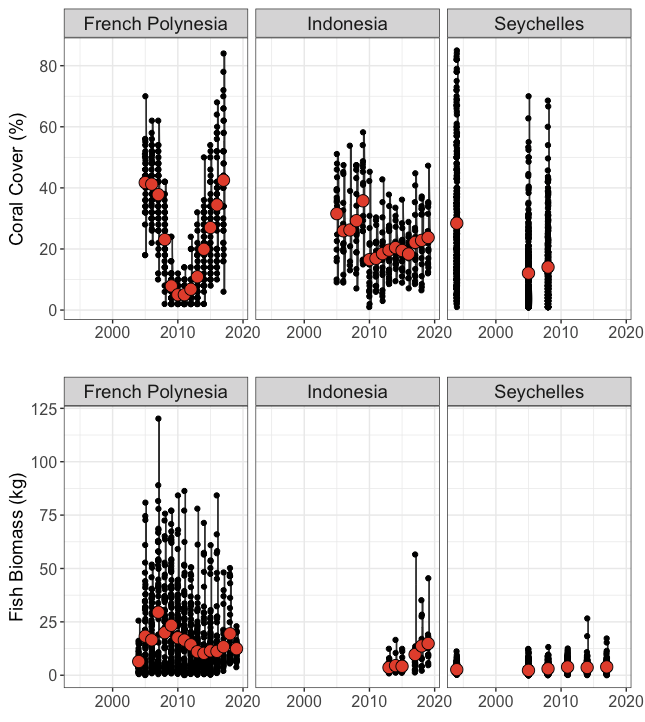

**Figure S5.** Trends in coral cover across time for the 38 time series belonging to the three regions (French Polynesia, Indonesia and Seychelles)

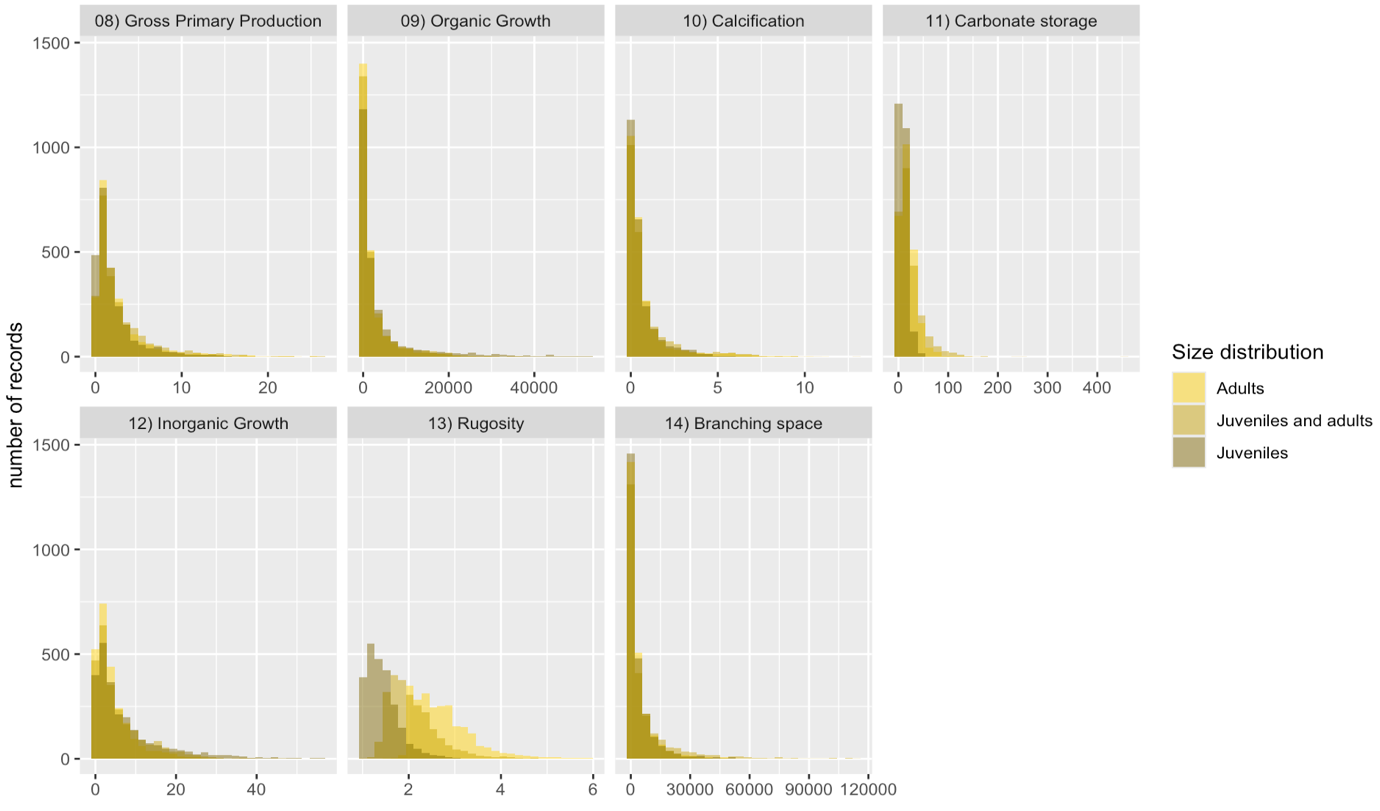

**Figure S6.** Sensitivity analysis for the different size structures according to the 7 benthic functions defined in the main manuscript.

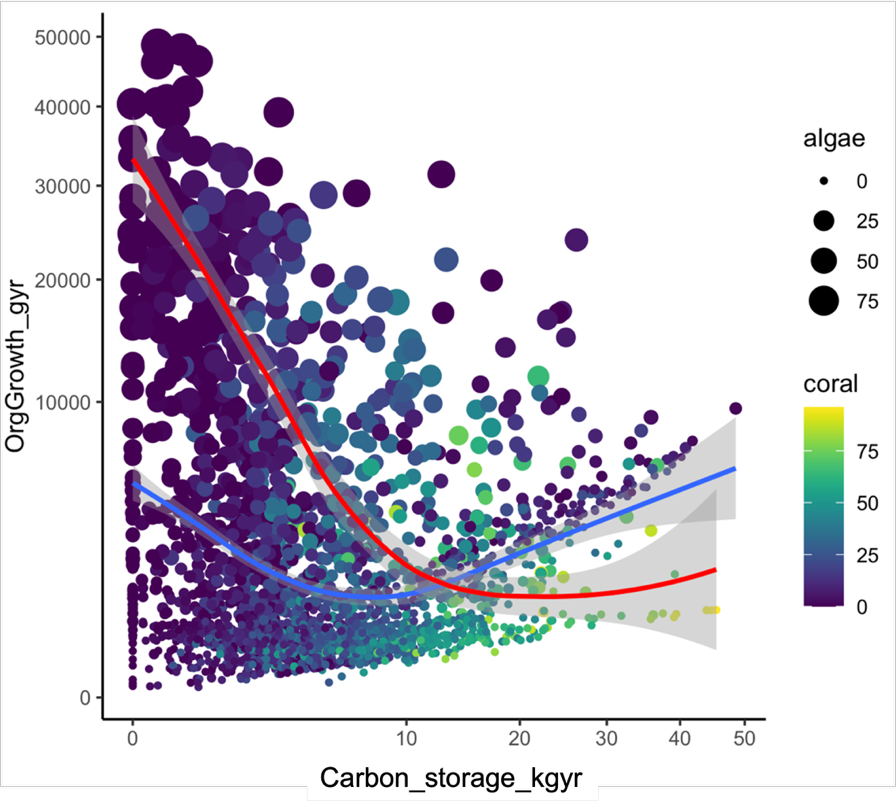

**Figure S7.** Relationship between organic production and carbonate storage on reefs examined in the study. Reefs (points) are colored by % coral cover and the size is proportional to % fleshy algae cover. While we expect a trade-off between these two functions, it does not emerge because of reefs with low coral and algal cover (blue line). When we consider reefs with >40% combined cover of coral and algae, the trade-off emerges (red line).

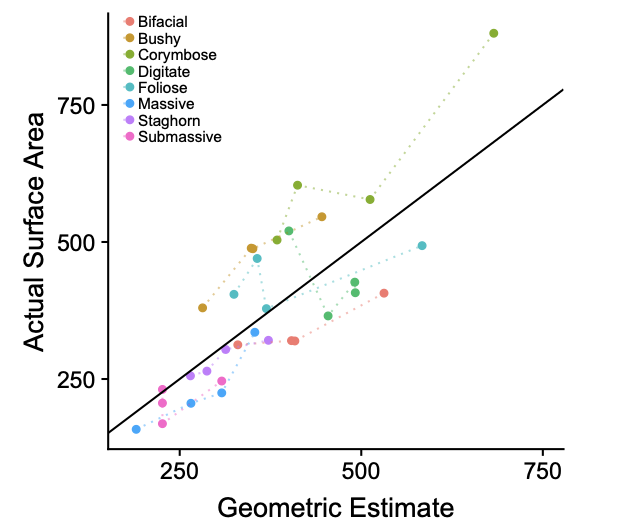

**Figure S8.** Test of the effectiveness of geometric estimates. The y-axis is surface area derived from a 3D scan. The x-axis is the estimate of surface area based on the equations in table S4.

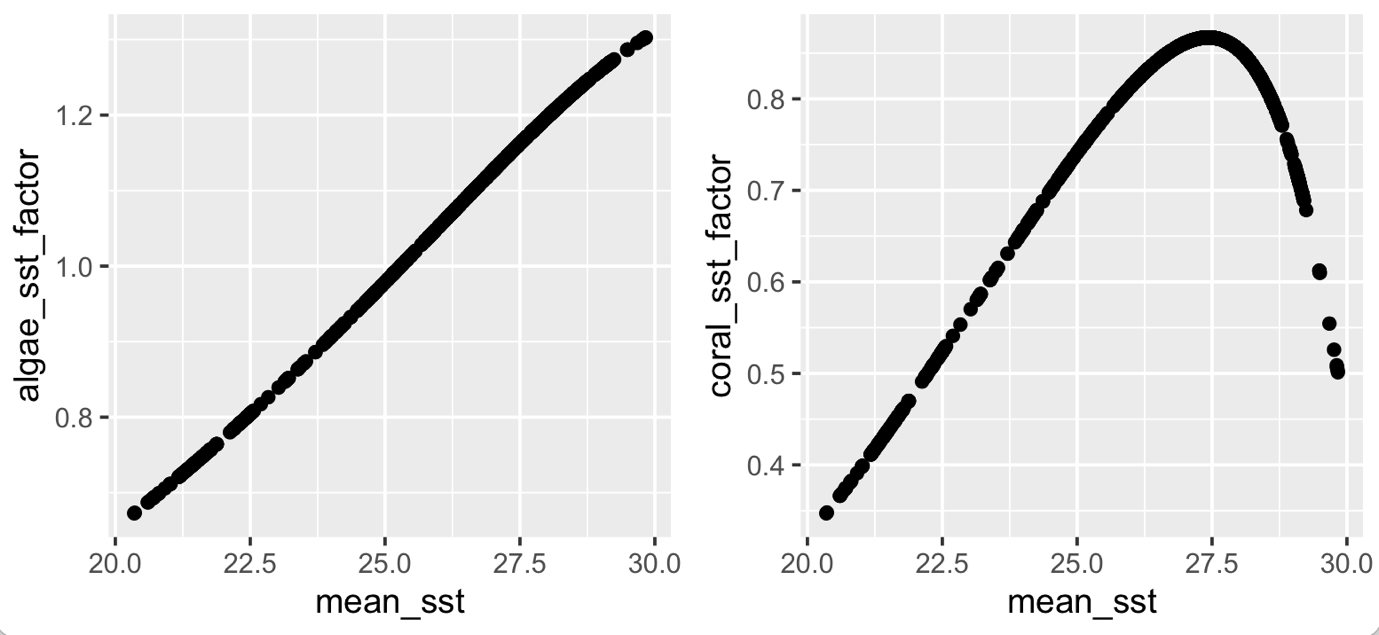

**Figure S9.** Temperature corrections factors used for corals and algae.

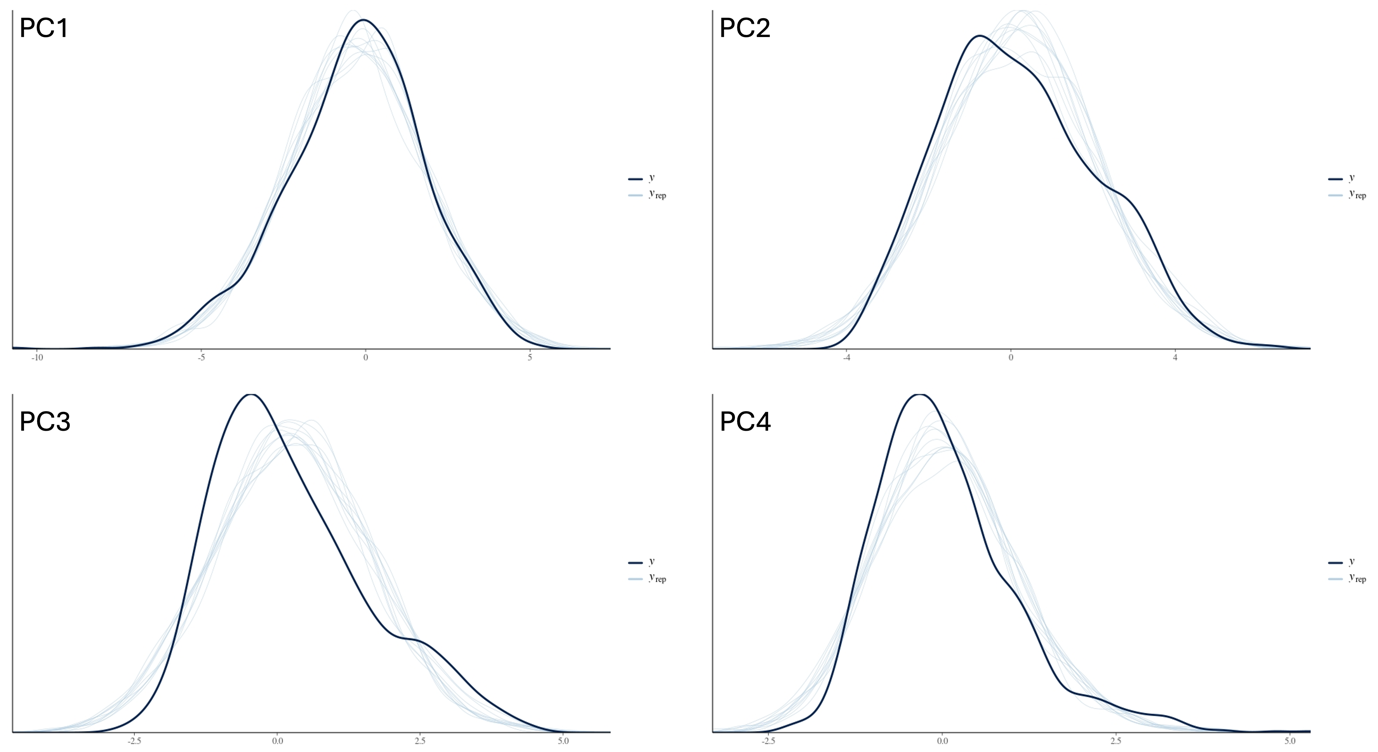

**Figure S10.** Posterior predictive check for the hierarchical bayesian models exploring the relationship between each Principal Component of the functional spectrum, environmental variables, DHW, Human Gravity and the presence of no-take MPAs

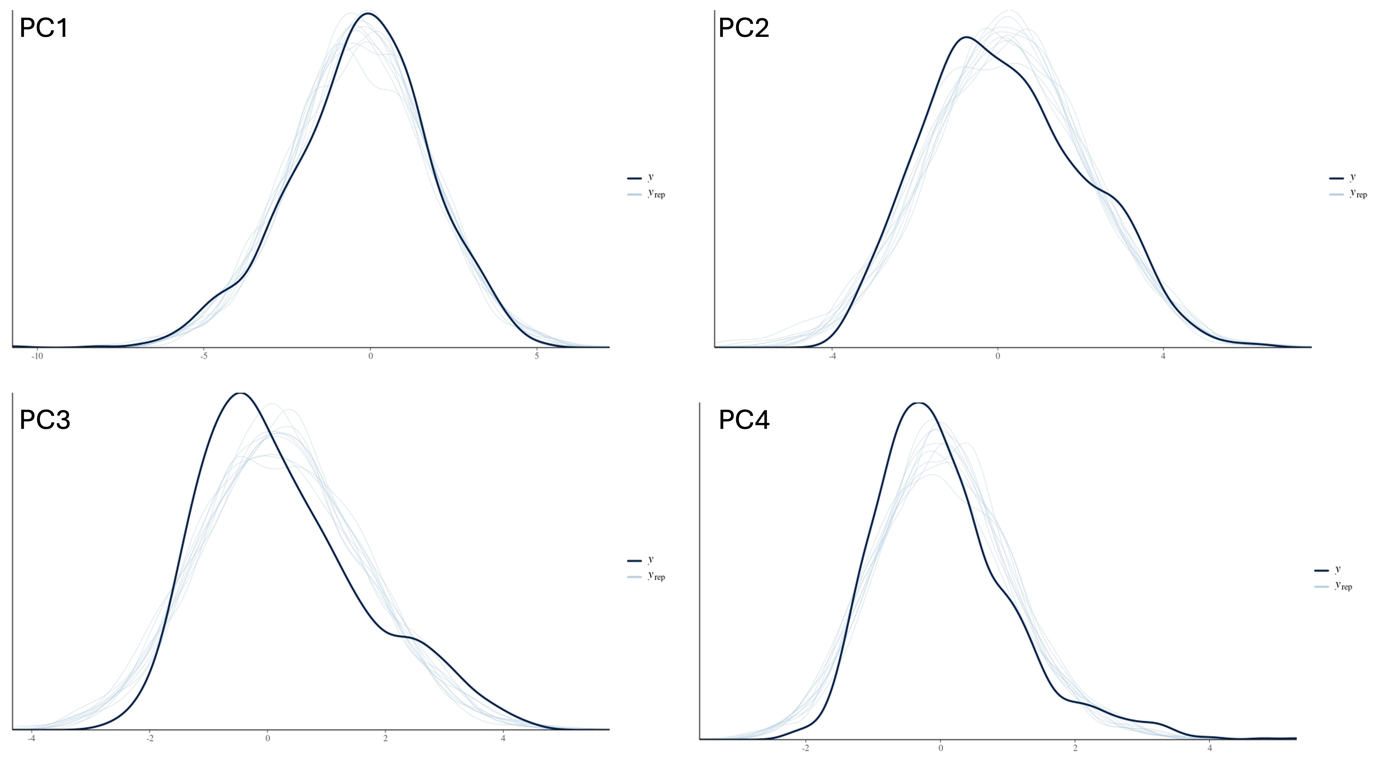

**Figure S11.** Posterior predictive check for the hierarchical bayesian models exploring the relationship between each Principal Component of the functional spectrum, DHW, Human Gravity and the presence of no-take MPAs (i.e. without environmental variables as explanatory variables).

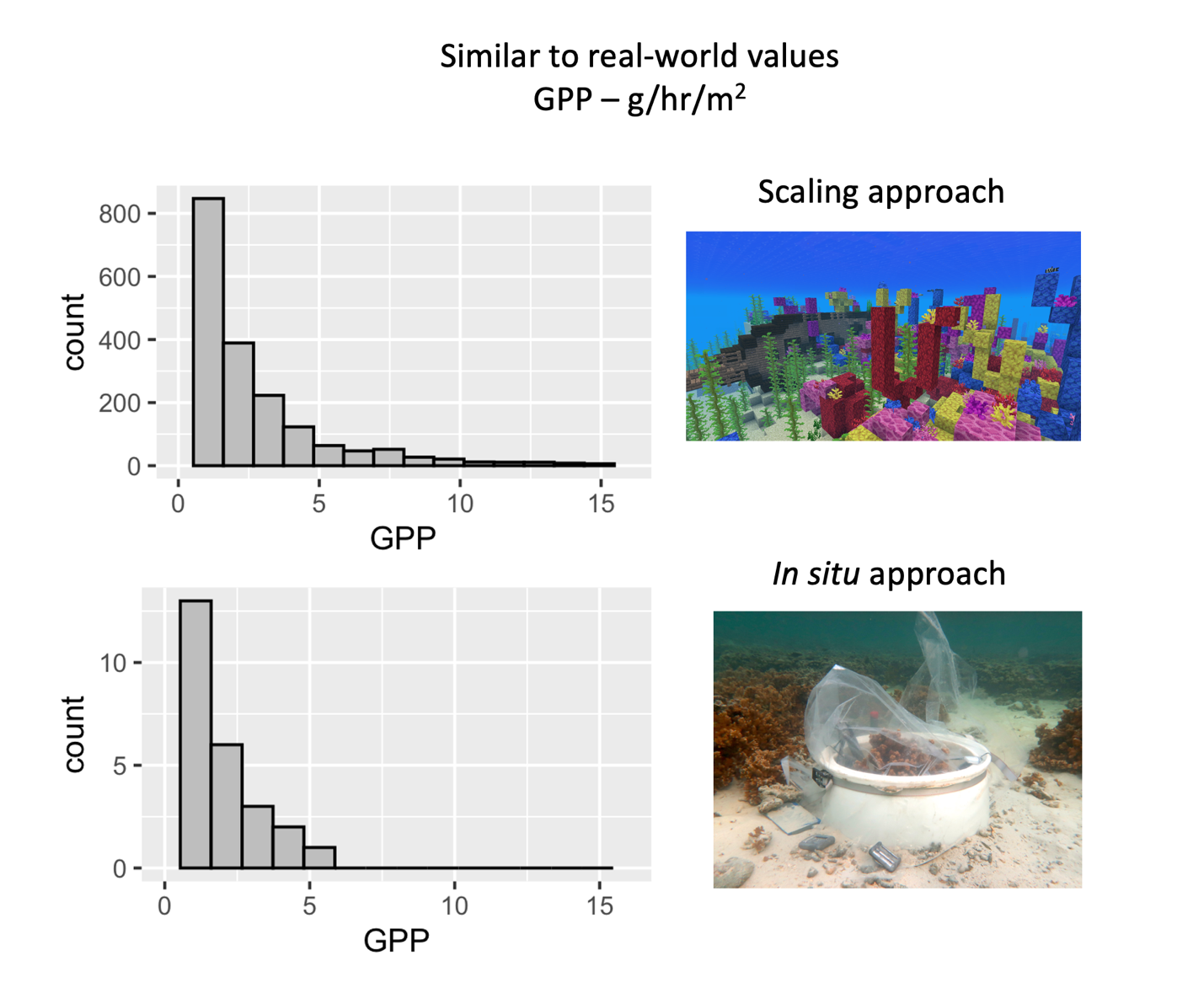

Figure S12. Frequency plot of primary productivity values derived from the scaling approach used here and *in situ* estimates of productivity derived from metabolic chambers.

**Table S1**. List of sites used for the temporal analyses, their geographic position and year of data collections

| **Region** | **Site** | **1994** | **2004** | **2005** | **2006** | **2007** | **2008** | **2009** | **2010** | **2011** | **2012** | **2013** | **2014** | **2015** | **2016** | **2017** | **2018** | **2019** |
| --- | --- | --- | --- | --- | --- | --- | --- | --- | --- | --- | --- | --- | --- | --- | --- | --- | --- | --- |
| Indonesia | B3 |  |  | x | x |  | x | x | x | x | x | x | x | x | x | x | x | x |
| Indonesia | KAL |  |  | x | x |  | x | x | x | x | x | x | x | x | x | x | x | x |
| Indonesia | KDS |  |  | x | x |  | x | x | x | x | x | x | x | x | x | x | x | x |
| Indonesia | PK |  |  | x | x |  | x | x | x | x | x | x | x | x | x | x | x | x |
| Indonesia | R1 |  |  | x | x |  | x | x | x | x | x | x | x | x | x | x | x | x |
| Indonesia | SAM |  |  | x | x |  | x | x | x | x | x | x | x | x | x | x | x | x |
| French Polynesia | Afareaitu |  | x | x | x | x | x | x | x | x | x | x | x | x | x | x |  |  |
| French Polynesia | Aroa |  | x | x | x | x | x | x | x | x | x | x | x | x | x | x |  |  |
| French Polynesia | Entre2baies |  | x | x | x | x | x | x | x | x | x | x | x | x | x | x |  |  |
| French Polynesia | Gendron |  | x | x | x | x | x | x | x | x | x | x | x | x | x | x |  |  |
| French Polynesia | Haapiti |  | x | x | x | x | x | x | x | x | x | x | x | x | x | x |  |  |
| French Polynesia | Maatea |  | x | x | x | x | x | x | x | x | x | x | x | x | x | x |  |  |
| French Polynesia | MotuAhi |  | x | x | x | x | x | x | x | x | x | x | x | x | x | x |  |  |
| French Polynesia | Nuarei |  |  | x | x | x | x | x | x | x | x | x | x | x | x | x |  |  |
| French Polynesia | Pihaena |  | x | x | x | x | x | x | x | x | x | x | x | x | x | x |  |  |
| French Polynesia | Taotaha |  | x | x | x | x | x | x | x | x | x | x | x | x | x | x |  |  |
| French Polynesia | Temae |  |  | x | x | x | x | x | x | x | x | x | x | x | x | x |  |  |
| French Polynesia | Tetaiuo |  | x | x | x | x | x | x | x | x | x | x | x | x | x | x |  |  |
| French Polynesia | Tiahura |  | x | x | x | x | x | x | x | x | x | x | x | x | x | x |  |  |
| Seychelles | CousinCarbonate | x |  | x |  |  | x |  |  | x |  |  | x |  |  |  |  |  |
| Seychelles | CousinGranite | x |  | x |  |  | x |  |  | x |  |  | x |  |  |  |  |  |
| Seychelles | CousinPatch | x |  | x |  |  | x |  |  | x |  |  | x |  |  |  |  |  |
| Seychelles | MaheECarbonate | x |  | x |  |  | x |  |  | x |  |  | x |  |  | x |  |  |
| Seychelles | MaheEGranite | x |  | x |  |  | x |  |  | x |  |  | x |  |  | x |  |  |
| Seychelles | MaheEPatch | x |  | x |  |  | x |  |  | x |  |  | x |  |  | x |  |  |
| Seychelles | MaheNWCarbonate | x |  | x |  |  | x |  |  | x |  |  | x |  |  | x |  |  |
| Seychelles | MaheNWGranite | x |  | x |  |  | x |  |  | x |  |  | x |  |  | x |  |  |
| Seychelles | MaheNWPatch | x |  | x |  |  | x |  |  | x |  |  | x |  |  | x |  |  |
| Seychelles | MaheWCarbonate | x |  | x |  |  | x |  |  | x |  |  | x |  |  | x |  |  |
| Seychelles | MaheWGranite | x |  | x |  |  | x |  |  | x |  |  | x |  |  | x |  |  |
| Seychelles | MaheWPatch | x |  | x |  |  | x |  |  | x |  |  | x |  |  | x |  |  |
| Seychelles | PraslinNECarbonate | x |  | x |  |  | x |  |  | x |  |  | x |  |  | x |  |  |
| Seychelles | PraslinNEGranite | x |  | x |  |  | x |  |  | x |  |  | x |  |  | x |  |  |
| Seychelles | PraslinNEPatch | x |  | x |  |  | x |  |  | x |  |  | x |  |  | x |  |  |
| Seychelles | PraslinSWCarbonate | x |  | x |  |  | x |  |  | x |  |  | x |  |  | x |  |  |
| Seychelles | PraslinSWGranite | x |  | x |  |  | x |  |  | x |  |  | x |  |  | x |  |  |
| Seychelles | PraslinSWPatch | x |  | x |  |  | x |  |  | x |  |  | x |  |  | x |  |  |
| Seychelles | SteAnneCarbonate | x |  | x |  |  | x |  |  | x |  |  | x |  |  | x |  |  |
| Seychelles | SteAnneGranite | x |  | x |  |  | x |  |  | x |  |  | x |  |  | x |  |  |
| Seychelles | SteAnnePatch | x |  | x |  |  | x |  |  | x |  |  | x |  |  | x |  |  |

**Table S2**. Literature search for trait values used in the estimation of benthic functions.

| **Source** | **doi** | **rls_group** |
| --- | --- | --- |
| Meyer et al | https://doi.org/10.1371/journal.pone.0133596 | Halimeda |
| Briggs and Carpenter 2019 | https://doi.org/10.1038/s43247-023-00766-w | coralline algae |
| Comeau et al. 2013 | https://doi.org/10.1038/s43247-023-00766-w | coralline algae |
| Comeau et al. 2014 JEMBE | https://doi.org/10.1038/s43247-023-00766-w | coralline algae |
| Comeau et al. 2019 Nat CC | https://doi.org/10.1038/s43247-023-00766-w | coralline algae |
| Tanaka et al. 2016 | https://doi.org/10.1038/s43247-023-00766-w | coralline algae |
| Johnson & Carpenter 2012 | http://dx.doi.org/10.1016/j.jembe.2012.08.005 | coralline algae |
| Meyer et al | https://doi.org/10.1371/journal.pone.0133596 | Halimeda |
| Cornwall et al. 2018 | https://doi.org/10.1038/s43247-023-00766-w | coralline algae |
| McWilliam et al 2018 | https://doi.org/10.1016/j.cub.2018.09.025 | coral |
| Tebbett & Bellwood 2021 | https://doi.org/10.1016/j.marenvres.2021.105311 | turf algae |
| Wefer 1980 | http://dx.doi.org/10.1038/285323a0 | Halimeda |
| Hudson 1985 | https://doi.org/10.1007/978-3-642-70355-3_20 | Halimeda |
| May-Lin et al 2013 | https://doi.org/10.1007/s10811-012-9963-5 | fleshy algae |
| Fischer and Martone 2014 | https://doi.org/10.1038/s43247-023-00766-w | coralline algae |
| Mackenzie & Agegian, Book Chapter | https://doi.org/10.1038/s43247-023-00766-w | coralline algae |
| Chan et al 2017 | https://doi.org/10.1002/2017GC006966 | coralline algae |
| Williams et al. | https://doi.org/10.1029/2020GL091499 | coralline algae |
| Drew 1983 | https://doi.org/10.1007/BF02395280 | Halimeda |
| Multer 1987 | https://doi.org/10.1007/BF00302014 | Halimeda |
| Mayukan & Panthrep 2019 | https://doi.org/10.1111/pre.12361 | Halimeda |
| Mayakun et al 2020 | https://doi.org/10.1017/ S0025315420001113 | Halimeda |
| Carpenter & Williams 2007 | https://doi.org/10.1007/s00227-006-0465-3 | turf algae |
| Vooren 1980 | https://doi.org/10.1016/0304-3770(81)90017-6 | turf algae |
| Jokiel & Morrissey 1986 Mar. Biol. | https://doi.org/10.1007/BF00397566 | fleshy algae |

**Table S3**. Geometric models used to project surface area and volume, adapted from McWilliam et al. 2018(*27*) with additional data from McWilliam et al. 2022(*15*)

| **RLS group** | **ctd_group** | **SA formula** | **Vol formula** |
| --- | --- | --- | --- |
| Branching | branching_open; branching_closed; hispidose | $\pi r_{c}^{2}(N_{b}\left( 2\pi r_{b}h_{b}+\pi r_{b}^{2} \right)$ | $\pi r_{c}^{2}(N_{b}\left( \pi r_{b}^{2}h_{b} \right))$ |
| Tabular | tables_or_plates | $\pi r_{c}^{2}(N_{b}\left( 2\pi r_{b}h_{b}+\pi r_{b}^{2} \right)$ | $\pi r_{c}^{2}(N_{b}\left( \pi r_{b}^{2}h_{b} \right))$ |
| Corymbose | corymbose; digitate | $\pi r_{c}^{2}(N_{b}\left( 2\pi r_{b}h_{b}+\pi r_{b}^{2} \right)$ | $\pi r_{c}^{2}(N_{b}\left( \pi r_{b}^{2}h_{b} \right))$ |
| Encrusting | encrusting | $\pi r_{c}^{2}$ | $\pi r_{c}^{2}h_{c}$ |
| Laminar | laminar; encrusting_uprights | $2\pi r_{c}\sqrt{r_{c}+h_{b}}$ | $h_{c}(\frac{1}{2}SA)$ |
| Hemispherical | massive; submassive; columnar | $2\pi r_{c}^{2}$ | $\frac{2}{3}\pi r_{c}^{3}$ |
| ahermatypic coral | NA | NA | NA |
| canopy forming macroalgae | NA | $\pi r_{c}^{2}(N_{b}\left( 2\pi r_{b}h_{b}+\pi r_{b}^{2} \right)$ | $\pi r_{c}^{2}(N_{b}\left( \pi r_{b}^{2}h_{b} \right))$ |
| seagrass | NA | NA | NA |
| understory macroalgae | NA | $\pi r_{c}^{2}(N_{b}\left( 2\pi r_{b}h_{b}+\pi r_{b}^{2} \right)$ | $\pi r_{c}^{2}(N_{b}\left( \pi r_{b}^{2}h_{b} \right))$ |
| fleshy algae | NA | $\pi r_{c}^{2}(N_{b}\left( 2\pi r_{b}h_{b}+\pi r_{b}^{2} \right)$ | $\pi r_{c}^{2}(N_{b}\left( \pi r_{b}^{2}h_{b} \right))$ |
| Halimeda | NA | $\pi r_{c}^{2}(N_{b}\left( 2\pi r_{b}h_{b}+\pi r_{b}^{2} \right)$ | $\pi r_{c}^{2}(N_{b}\left( \pi r_{b}^{2}h_{b} \right))$ |
| turf algae | NA | $\pi r_{c}^{2}$ | $\pi r_{c}^{2}h_{c}$ |
| slime | NA | NA | NA |
| coralline algae | NA | $\pi r_{c}^{2}$ | $\pi r_{c}^{2}h_{c}$ |

**Table S4.** Equations used to calculate 7 benthic functions.

| **Function** | **Formula** | **Units** |
| --- | --- | --- |
| Gross Photosynthesis | GPP (per cm^2^) * SA | g.hr^-1^ |
| Calcification | Calcification (per cm^2^) * SA | g.hr^-1^ |
| Carbonate storage | Volume * Skeletal density | kg |
| Inorganic production | [Vol_t1_ – Vol_t0_] * Skeletal density | kg.yr^-1^ |
| Organic Production | [Vol_t1_ – Vol_t0_] * Tissue biomass (for algae)  SA_t1_ – SA_t0_] * Tissue biomass (for coral) | g.yr^-1^ |
| Rugosity | Scaled up from Husband *et al.*, 2022 | unitless |
| Branch Space | Convex Hull (L * W * H) - Vol | cm^3^ |
